# The Biochemical Basis of Hormesis

**DOI:** 10.64898/2026.04.20.719646

**Authors:** Guillermo Cerrillo, Helena Vidakovic, David G. Míguez

## Abstract

Atypical dose-response curves, where the effect of a drug does not follow a monotonic shape, are unsuitable but common in drug development. Here, we develop a high-throughput computational screening to search for features that could induce this type of complex hormetic relation between changes in drug concentration and changes in drug effect. Our study suggests that all hormetic networks share a similar core structure: an incoherent bivalent network motif centered on the target of the drug. In addition, the probability of producing a biphasic dose response requires that one of the interactions operating in the saturated regime. This insight is applied to explain how hormesis exhibited during long-term rapamycin treatment arises directly from the structure of the network of interactions that form the mTOR-Pi3K-Akt signaling pathway. Overall, our study shows that drug hormesis can be explained and predicted, and should not be a cause for automatic rejection in the drug development pipeline.

## Introduction

Cells sense, process, and react to stimuli through an intricate network of interacting proteins. Disruptions in these signaling pathways due to inherited or *de novo* genetic alterations may affect the way cells translate these input stimuli into coordinated biological responses. At the tissue level, they can affect homeostasis and cell communication, and are at the core of many degenerative, autoimmune, or proliferative pathologies. Treatment strategies for these diseases with a clear molecular basis include monoclonal antibodies, gene therapy, and targeted small-molecule therapies. In this last category, small-molecule kinase inhibitors stand out as a powerful, efficient, and rapidly developing strategy, with around 94 of them currently approved by the FDA [1–2] (10 newly approved in 2025 and around 200 currently undergoing clinical trial validation). One of the main requirements for approval is a reliable and predictable relation between dosage and activity. In other words, a dose-response curve has to follow a typical sigmoidal shape as the concentration increases (below toxic levels), with a well-defined IC50 (the inflection point where the drug reaches 50% efficiency).

In this context, a nontypical dose-response curve constitutes a major red flag, often interpreted as an indication of potential problems or side effects. For instance, drugs with the highest efficiency at a narrow range of concentrations, or even drugs that act as inhibitors at lower doses and activators at higher doses. In the field of drug development, these types of complex dose-responses are commonly referred to as *hormetic* or *biphasic*.

A systematic analysis of peer-reviewed literature identified around 9000 dose–response relationships that can be classified as hormetic (higher frequency of appearance than the typical sigmoidal shape) [3–4]. Compounds as widely used as aspirin (reduces recurrent cardiovascular events at intermediate doses, but not at low or high doses) [5], heparin (anti-inflammatory at low doses, pro-inflammatory at high doses) [6], progesterone (decreases plasminogen activator inhibitor type 1 antigen only at intermediate doses) [7]. In addition, nicotinamide (a common neuroprotective agent) [8], metformin (treatment for Type 2 diabetes) [9], several antifungal, antibacterial, antiviral, and antitumour agents (such as curcumin [10] or ATN-161 Inhibitor [11], show a clear biphasic response.

Arguably, the most widely reported hormetic drug is rapamycin [12–14], an FDA-approved macrolide antibiotic used clinically to induce immunosuppression in organ-transplant patients, vascular anomalies, as well as certain autoimmune conditions and cancers [15]. In the last decade, rapamycin has been shown to have a remarkable potential to increase maximum life span in yeast [16], worms, flies, and mice [17]. In humans, it has been shown to improve multiple physiological parameters associated with aging [17]. Rapamycin and its derivatives (rapalogs) work basically as mTOR inhibitors, a protein [18–19] involved in many aspects of the aging process [20]. The hormetic characteristics of rapamycin have been widely documented [21–22], with a positive effect at intermediate concentrations, and a detrimental effect at high doses, commonly attributed to toxicity [23–24].

A potential source of complex drug responses is the fact that targets are part of signaling networks, and these can be highly nonlinear in their structure and in the nature of their interactions. Signaling pathways often contain recurrent regulatory motifs with an important role in translating an external stimulus into an appropriate downstream response. The effect of these nonlinear network motifs has been extensively studied and characterized both theoretically and experimentally, and has been shown to influence robustness, adaptation, stability, and dynamics. In this context, negative feedback is known to induce desensitization [25–26], positive feedback can induce bistability, and a feedforward loop induces fold change dependence [27]. We have previously shown that certain network motifs can induce a previously unknown form of inverse hysteresis, producing a bistable reversible desensitization to drug treatment [28].

Interestingly, none of the known nonlinear network motifs or any combination of them has been associated with a disruption in the monotonic shape of the dose-response curve. In this direction, the molecular mechanisms that can induce drug hormesis are not known. To investigate this, we designed a high-throughput computational screening to answer the following questions: Are there specific signaling pathway topologies that can induce biphasic dose-response curves? What are the required ingredients to induce hormesis? Can we explain the hormesis reported for key compounds based, such as rapamycin?

Our study demonstrates that network topologies where the target of inhibition is part of an incoherent bivalent motif are prone to exhibit biphasic dose-response curves. In addition, if one of the bivalent interactions of the target operates in a saturated regime (high affinity towards its substrate), the probability of hormesis is highly increased. Finally, we apply this insight to explain the hormesis of rapamycin, which arises from the specific topology of the mTOR-PI3K signaling cascade at long exposures.

## Materials And Methods

### Main approach and design

Our approach uses a high-throughput computational approach to screen all potential network topologies in a conceptual coarse-grained model of a signaling cascade. The model is centered around a small molecule inhibitor that binds to its *Target* molecule. Molecular interactions above this *Target* are condensed as part of a single *Input* node, while downstream processes are condensed as an *Output* node. A scheme of the approach is illustrated in Fig 1. The signaling starts at the *Input* node, which receives a constant activation signal. The dynamics of the downstream cascade of activation and deactivation events is computed by solving the corresponding set of coupled ODEs. The value of the *Output* node at steady state is recorded, and the entire process is then repeated for increasing concentrations of the inhibitor, constructing the corresponding dose-response curve. Dose-response curves with a difference between their maximum and minimum values lower than 0.01 are labeled as *insensitive* to the drug treatment. The remaining dose-response curves are classified as *Hormetic* if there is a change in the sign of the local slope. Topologies with biphasic curves higher than 1% are classified as “hormetic” and are stored for further classification.

**Figure 1:**
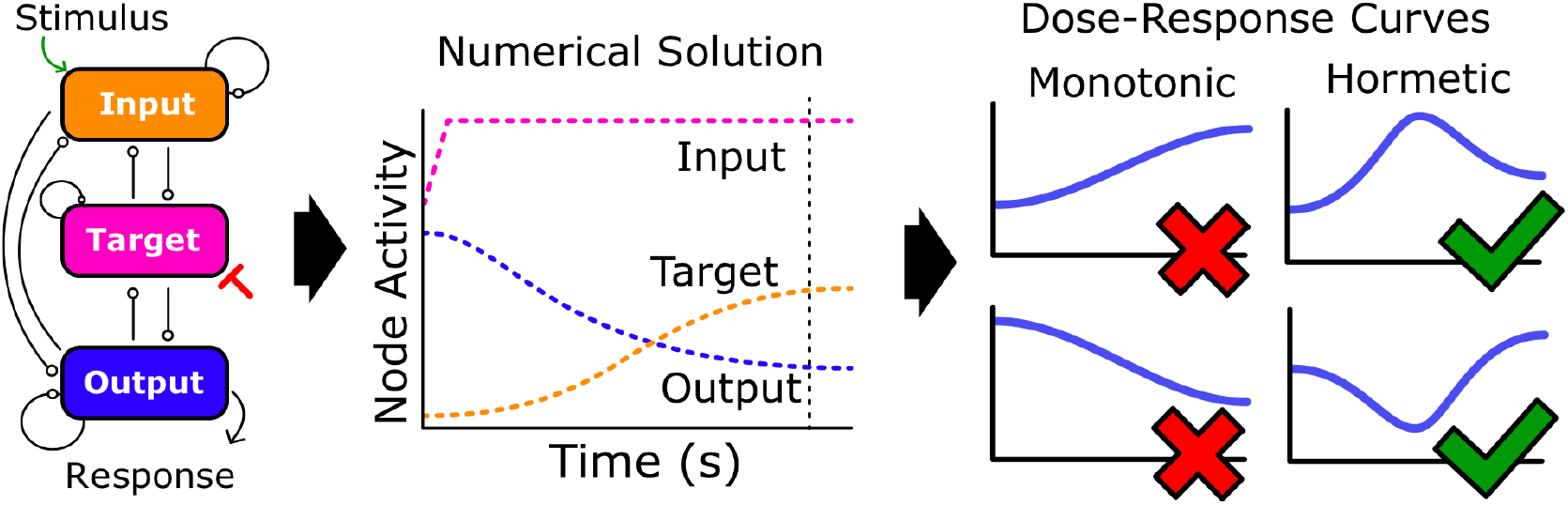
Scheme of the approach used. (left panel) All potential interaction schemes between three nodes are tested using a high-throughput computational approach. The Input node is constantly stimulated, the Target node is inhibited by our inhibitor, and the activity is monitored based on the Output node. A total of 5000 numerical simulations for each potential network topology (to sample the entire parameter space) are computed until a steady state is reached (central panel). The value of the Output at steady state (vertical dashed line) is plotted for each inhibitor concentration as a dose-response curve. These curves are evaluated and classified as monotonic or hormetic (right panel) based on changes in the sign of the derivative. The details of the framework are explained in the Methods section.

To sample all the parameter space, this process is repeated 5000 times using different values for the kinetic parameters, resulting in 5000 different dose-response curves per topology. Finally, all potential sets of interaction topologies between *Input, Target*, and *Output* nodes are evaluated. Since input and output nodes represent multiple interactions (above and below the *Target* protein), feedback regulation (positive and negative) is allowed in these nodes. For simplicity, the *Target* molecule is assumed not to directly activate or deactivate itself, only indirectly via the input and output nodes. In addition, interaction topologies where the *Input, Target*, or *Output* nodes are disconnected are not evaluated, and only one interaction between each pair of nodes is allowed. These conditions reduce the set of 3^3×3^ = 19683 potential network topologies to only 5346.

### Model Equations and Parameter Sampling

The kinetic equations of the model are defined as three coupled ODEs where interactions between *Input, Target*, and *Output* nodes are computed following a Michaelis-Menten approach of activation-deactivation, in its general form:

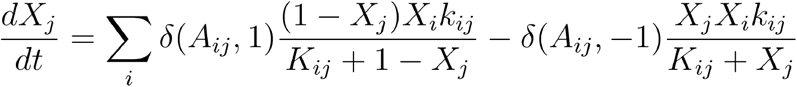

where *X*_*j*_ and *X*_*i*_ (i,j=1…3) correspond to the active form of *Input, Target*, and *Output* nodes, (0< *X*_*j*_ *<1*, with *X*_*j*_ *= 1* corresponding to full activation). Parameter *k*_*ij*_ corresponds to the kinetic constant of the interaction of species *i* activating species *j*, and *K*_*ij*_ is the Michaelis-Menten constant of this specific interaction.

Computationally, this process is implemented by transcribing all potential topologies as adjacency matrices *A* of 3×3, where the element *A*_*ij*_ = 1 if there is direct activation of node *j* mediated by node *i, A*_*ij*_ = −1 if there is direct inhibition of node *j* mediated by node *i*, and *A*_*ij*_ = 0 if there is no interaction from node *i* to node *j*. For instance, a value of *A*_*13*_ = −1 means that the *Input* node is directly inhibiting the *Output* node, while *A*_*33*_ = 1 means that the *Output* is directly activating itself. Since there is no direct feedback in the *Target*, the value of *A*_*22*_ = 0. The δ symbol represents the Kronecker delta function, which can take values of 1 or 0 depending on the topology being evaluated: δ(A_ij_,1) = 1 only if node i activates node j, δ(A_ij_,-1) = 1 only if node i inhibits node j.

To avoid the trivial steady state where a node is fully activated if it only receives activating interactions, the software adds a background deactivation interaction as a balance (otherwise). Similarly, if a given node only receives deactivating interactions, the software adds a background activation interaction. Finally, the upstream activation of the network is included as constant activation of the *Input* node:

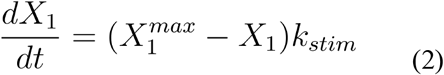

The inhibition of the *Target* node is implemented by modulating k_21_ and k_23_ (i.e., all interactions where the *Target* is acting) by:

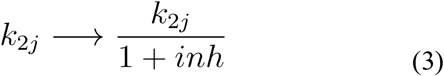

where *inh* is the concentration of the inhibitor, which is maintained constant throughout each simulation. For each set, 30 inhibitor concentrations were tested, from *inh =* 0.05 to 1000, distributed on a logarithmic scale. Parameter values are summarized in Table 1.

| Parameter | Value | Description |
| --- | --- | --- |
| $K_{ij}$ | Random between 0.001 and 100 | Michaelis-Menten Constant |
| $k_{ij}$ | Random between 0.1 and 10 | Kinetic Constant |
| $X_i, X_j$ | computed | amount of active nodes |
| $X_1^0$ | fixed at 0.0 | Initial input activation |
| $X_2^0$ | fixed at 0.5 | Initial target activation |
| $X_3^0$ | fixed at 0.5 | Initial output activation |
| $inh$ | 30 values from 0.05 to 100 | Inhibitor concentration |
| $\delta(A_{ij}, 1)$ | 1 if $A_{ij}=1$, 0 if $A_{ij} \neq 1$ | Delta Kroneker |
| $\delta(A_{ij}, -1)$ | 1 if $A_{ij}=-1$, 0 if $A_{ij} \neq -1$ | Delta Kroneker |
| $X_1^{\max}$ | 0.5 | Activation by stimulus |
| $k_{\text{stim}}$ | 0.1 | kinetic constant of stimulus |

### Mass action model of the mTOR-Pi3K pathway and the effect of Rapamycin

The mathematical model of the *Pi3K-mTOR* pathway and the interaction of rapamycin is implemented as a set of coupled ODEs derived using full mass action kinetics. A scheme of the interactions included in the model is illustrated in Figure 5A, based on the following interactions (marked with red numbers).

- 1. The receptor IRS upstream of the pathway is activated by binding reversibly to Insulin [29]
- 2. IRS is inactivated by Phosphorylation by the active form of S6K1/2 [14, 30].
- 3. Active IRS promotes the activation of Pi3K by phosphorylation [31].
- 4. mTOR can bind to raptor to form the complex mTORC1, or to rictor to form mTORC2 [13].
- 5. mTORC1 activates S6K1/2 by phosphorylation [32].
- 6. Rapamycin binds specifically to mTORC1 in a reversible manner [12].
- 7. Active Pi3K activates mTORC2 by phosphorylation [33].
- 8. Active Pi3K activates IRS1 indirectly via AKT, forming a positive feedback loop [34–35].
- 9. Active S6K1/2 deactivates mTORC2 [36].
- 10. PTEn is the main phosphatase that indirectly inactivates Pi3K [37].
- 11. PP2A is the main phosphatase that counteracts the effect of mTORC1 in the activation of S6K1/2 [38–39].

The corresponding differential equations are:

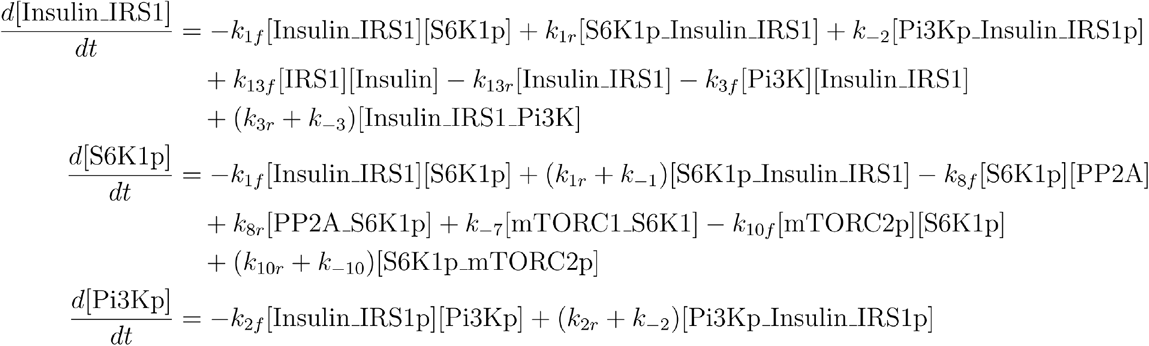

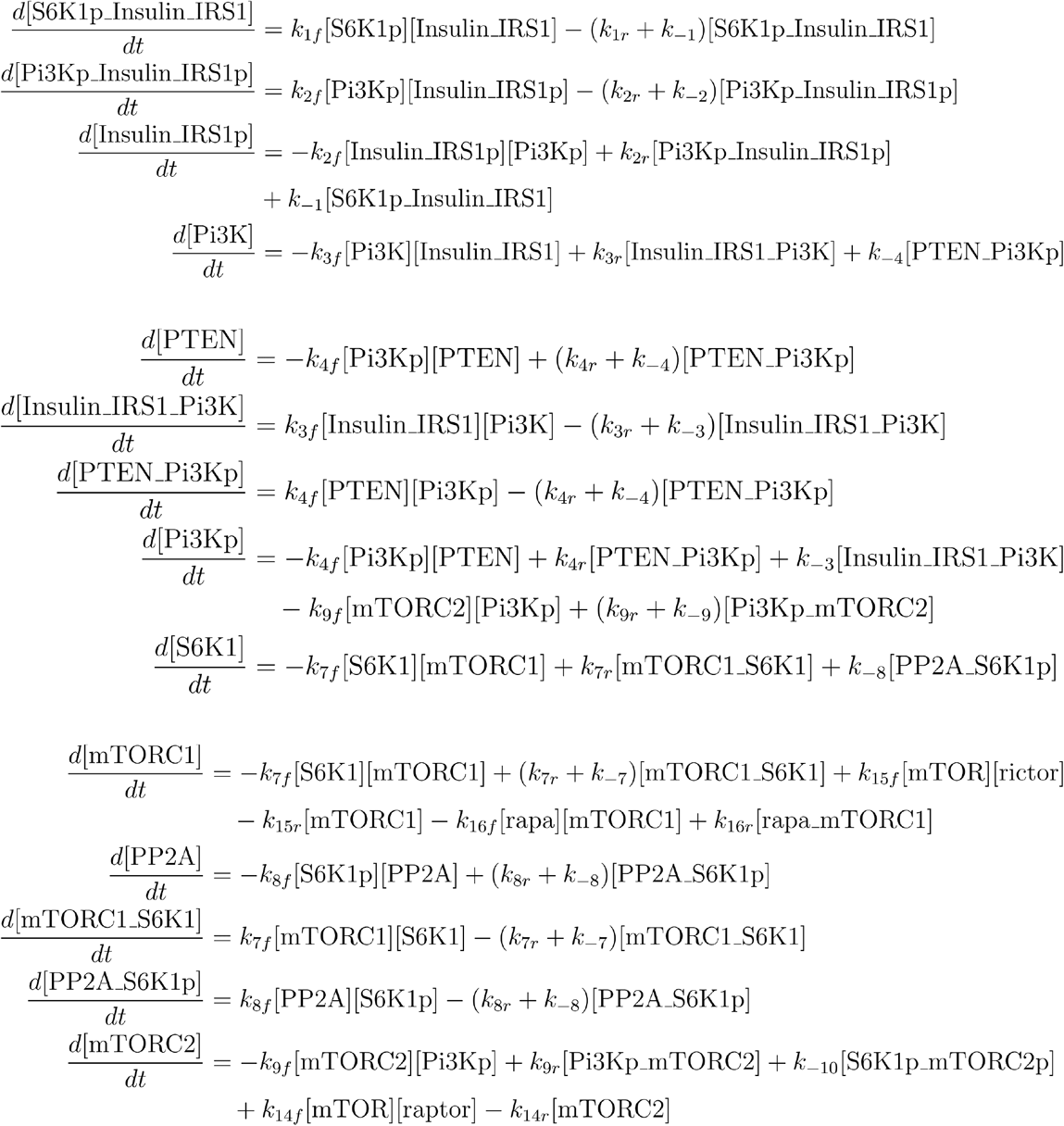

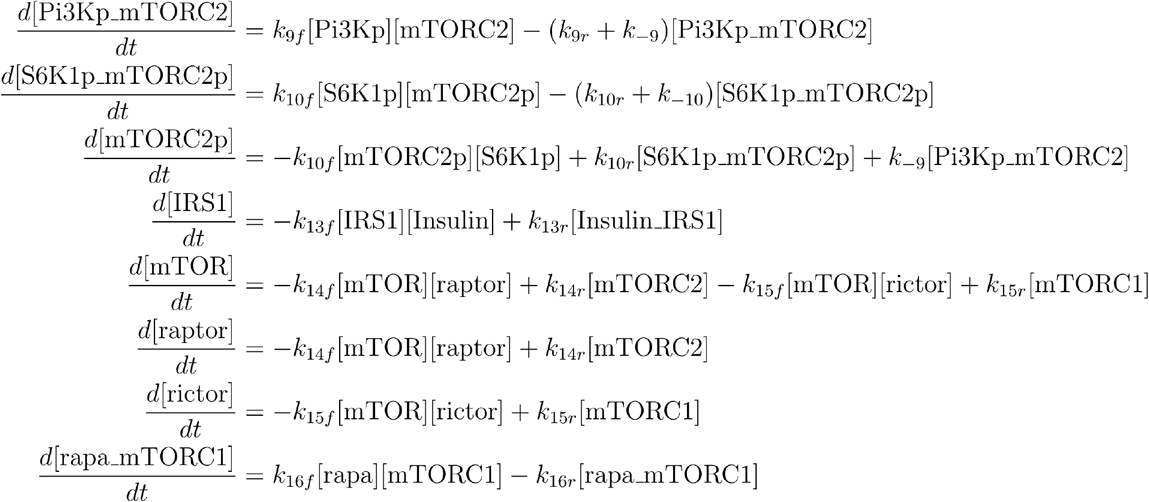

### Computer requirements, algorithms, and code

The approach resulted in a total of 5346 (topologies) × 5000 (parameter sets) × 30 (inhibitor concentrations) ≈ 8×10^8^ simulations, performed on a desktop computer running Windows 11 Pro, with an Intel Core i7-13700 (24 CPUs ~2.1 GHz) and 32 GB of RAM. Code is written in Julia 1.11.4. (available in https://github.com/GuillermoCerrillo/TFM_Hormesis). The model is written in Julia [40] and solved using a Runge-Kutta algorithm implemented in the package *DifferentialEquations* [41]. The solver *AutoTsit5* is used by default. If stiff changes are detected for a particular set of parameters, the program switches to the Rosenbrock23 solver (relative tolerance was set to 1e-8, absolute tolerance was set to 1e-10). Simulations are computed for a minimum of 500 time steps. If equilibrium is not reached at this point (set as a change of 1e-5 between consecutive time points), the simulation continues until equilibrium is reached. If the simulation reaches 2500 time steps, it is discarded from the analysis. If this happens a total of 10 times per set, or in 4 consecutive points of the dose-response curve, the set is discarded.

Sampling of the whole parameter space was performed by generating 5000 random sets of *k* and *K* parameters (Latin Hypercube Sampling, from 0.1 to 10 for *k*_*ij*_, 0.001 to 100 for *K*_*ij*_).

## Results

### A large number of topologies exhibit biphasic dose-response curves

A first general characterization of the overall probabilities of hormesis is summarized in Figure 2. Our analysis shows that more than 500 of the possible topologies tested show clear non-monotonic dose-response curves (Figure 2A). When organized by the number of interactions, we can clearly see that the highest number of topologies that are capable of exhibiting hormesis contain 6 or 7 links (Figure 2B), suggesting that the probability of hormesis increases as the network becomes more complex. Figure 2C shows the same data but normalized by the number of possible topologies per number of links (there are many more possible networks of 5 links than of 8 links, for instance). The increase in the number of hormetic topologies as the number of active links increases indicates that the probability of exhibiting hormesis depends on the density and complexity of the network.

**Figure 2:**
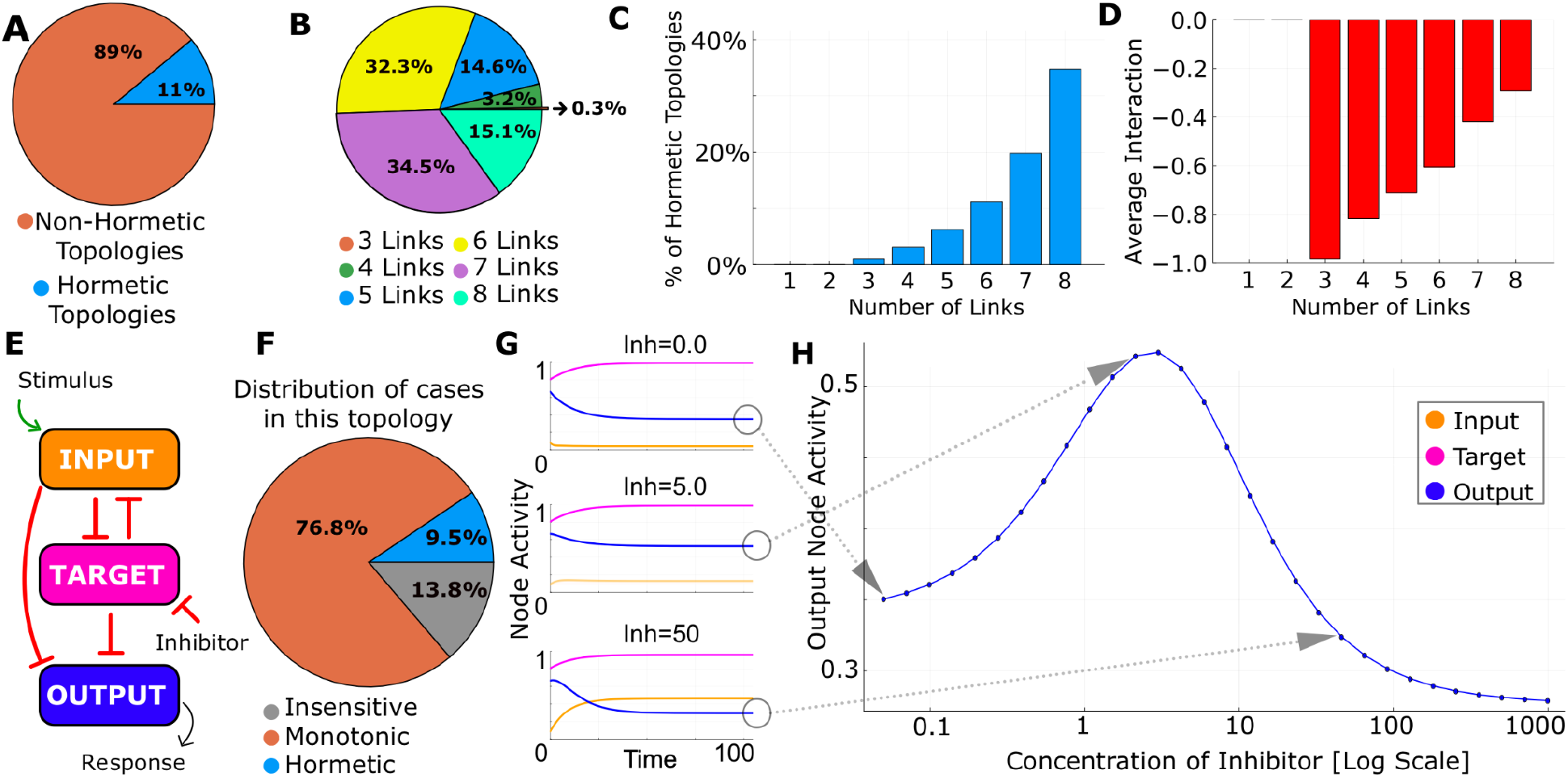
Overall Characterization of hormetic network topologies. (A) Percentage of network topologies capable of producing a nonmonotonic dose-response curve. (B) Distribution of topologies capable of producing hormesis by number of links. (C) Percentage of topologies for a given number of links that can produce hormesis. (D) Average sign of the interactions clustered by the number of links in the network. (E) Scheme of the network topology with the highest percentage of Hormetic drug-response curves. (F) Distribution of hormesis versus monotonic drug responses in the previous network. (G) Dynamics of the three nodes for low, medium, and high inhibitor concentration, for a given set of parameter values. (H) Example of a hormetic Dose-Response curve for the previous network topology.

Next, we focus on investigating what type of interactions induce hormesis with the highest probability. To do this, we plot in Figure 2D the sum of all components of the adjacency matrices for all hormetic topologies. In this representation, a negative value corresponds to a tendency for deactivating or repressing interactions on average, and a positive value corresponds to an average of activating interactions. This analysis shows a clear tendency towards interactions of the repressing type, with this trend reducing towards a balance between positive and negative interactions as the networks grow in complexity.

Figure 2D focuses on the topology that produced the highest number of biphasic dose-response curves in our analysis (composed of four repressing interactions). Out of the 5000 conditions tested, a total of 475 dose response curves can be classified as nonmonotonic (more than 11% of all parameter sets that show responsiveness to inhibition, Figure 2E). To understand how the biphasic response emerges in this particular topology, we plot three numerical simulations of this topology for low, medium, and high concentrations of inhibitor (the rest of the parameters are maintained). We can see that in the three situations, the *Target* (pink line) remains constant at high values, while the concentration of *Input* increases as the inhibitor increases (orange line) because the *Target* is negatively regulating the *Input* node. This increase in the concentration of the *Input* increases the repression of the *Output* mediated by the *Input* node. Interestingly, the effect of inhibition via the *Input* node seems prominent at high inhibitor concentrations, while the direct repression of the *Output* mediated by the *Target* seems to be the dominant interaction at lower concentrations of inhibitor. Overall, the biphasic response in the *Output* node in this particular topology seems to occur due to a balance between activation and repression, taking preference at different concentrations of inhibitor. This balance generates a dose-response curve plotted in Figure 2E, with a clear inverted U shape.

In conclusion, this general characterization shows that hormesis is a common feature that can arise even in these types of smaller network motifs, and that it is clearly linked to network complexity and favored in networks with a high number of repressing interactions on average. A detailed analysis of the biphasic response in one network shows that the combination of two interactions modulating the activity of the *Target*, one acting at low inhibitor and the other more dominant when the inhibitor is high, seems to be underlying the biphasic dose-response curve.

### Hormesis arises from four minimal topologies with a similar core structure

Next, we proceed to study whether there are core interactions or interactions common to all topologies classified by our analysis as hormetic. To do that, we start with the largest networks (8-links) and compare their percentage of hormesis with their parental networks (i.e., networks of 7 links that only differ by a single interaction). If this extra link does not increase the percentage of hormesis, we assume that the core architecture required for hormesis is the same in both networks and that the extra link does not positively influence their hormetic nature. If this is the case, the 8-link topology is discarded from the analysis. This is then repeated between networks of 7 and 6 links, between 6 and 5, then between 5 and 4, and finally between 4 and 3 links, allowing us to filter out all redundant network topologies and obtain a subset of all topologies where each link has a net positive effect in the ability to generate biphasic dose-responses. This new set contains only 16 basic topologies, distributed by link density as shown in Figure 3A.

**Figure 3:**
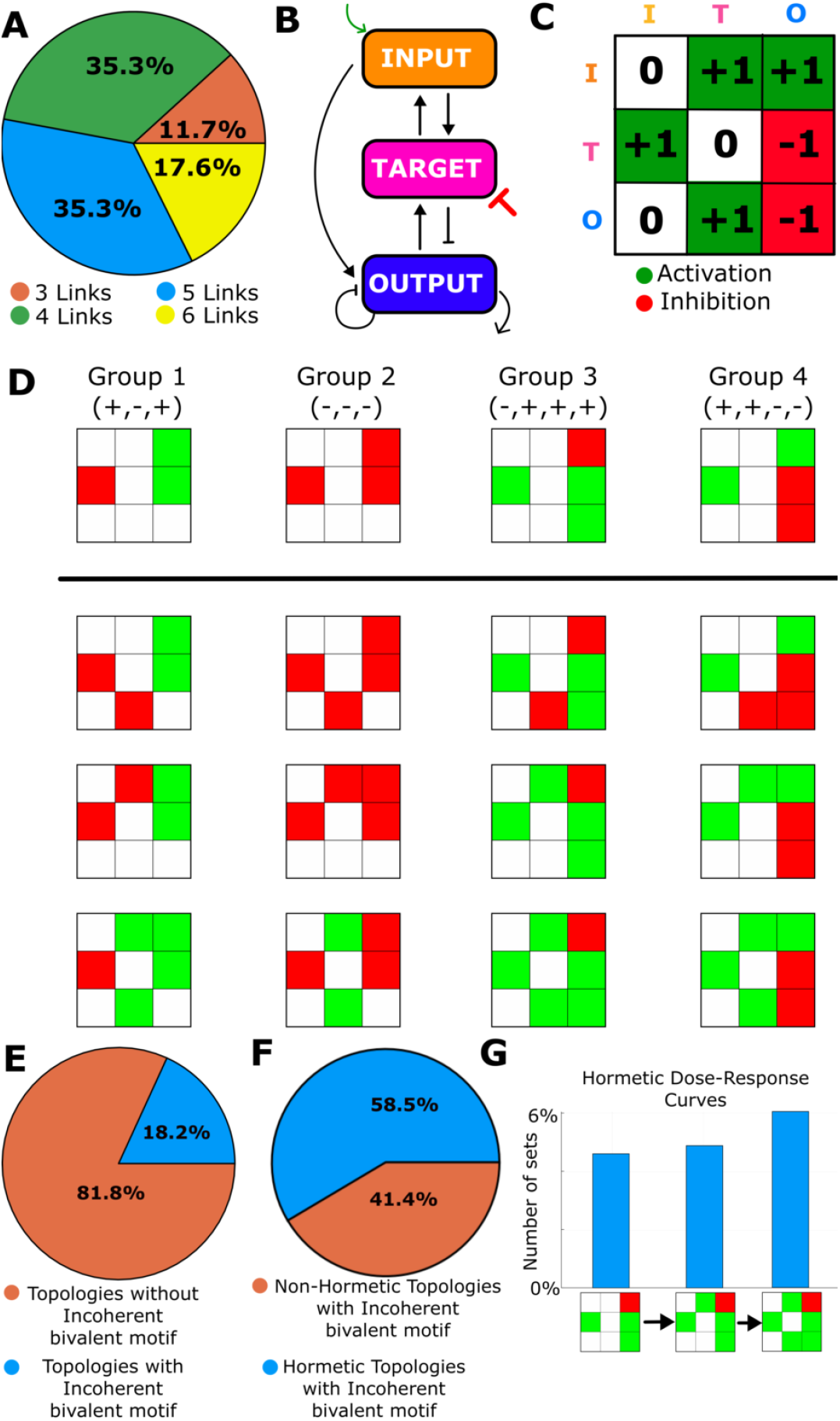
Hormesis appears in networks within four core incoherent bivalent motifs. (A) Distribution of network topologies capable of producing nonmonotonic dose-response curves after redundant topologies are filtered. (B-C) Example of a hormetic topology and its illustration as a color diagram. (D) Organization of the 16 Hormetic network topologies of three nodes based on the presence of one of the four core network motifs. (E) Percentage of topologies that have an Incoherent Bivalent Motif centered in the Target node. (F) Percentage of topologies with an Incoherent Bivalent Motif centered in the Target node that produce hormetic dose-response curves. (G) Increase in percentage of Hormetic cases after adding extra links to a core topology.

Next, we proceed to cluster them in families based on common minimal topological motifs. To do that, we generate a visual representation of the adjacency matrix for each topology as a 3X3 square diagram where links are color-coded by sign (red corresponds to negative interaction, green corresponds to positive interaction, and white corresponds to no interaction). An illustration of this scheme is shown in Figure 3B-C, where a hormetic topology is represented by the diagram on the right (rows represent links starting from *Input, Target*, and *Output*, and columns correspond to links arriving at the *Input, Target*, and *Output*). This representation is then used to illustrate the 16 core topologies as four main families (columns in Figure 3D), each one sharing a core motif (first row). The main common characteristic in all four families is that all have a link from *Target* to *Input*, i.e, pointing upstream to the signal flow (from stimulus to *Output*). This link can be either inhibitory (Groups 1 and 2) or activatory (Groups 3 and 4). In addition, all networks have an additional interaction from *Target* to *Output*. This dual role of the *Target* found in all core topologies is commonly referred to as a bivalent motif. In our particular case, one of the interactions needs to go upstream, while the other points downstream in the sense of the signaling induced by the stimulus.

Focusing on the other interactions, if the link from *Target* to *Input* is inhibitory, the other two links in the network have to be either positive (Group 1) or negative (Group 2). On the other hand, if the *Target* activates the *Input*, the network needs two non-feedback links of different sign, combined with a feedback in the *Output* node that can be positive (Group 3) or negative (Group 4). In other words, the requirement for a network to exhibit hormesis is that the two links of the bivalent node have net opposite (direct or indirect) effects in the *Target*, in the form of an incoherent bivalent motif. This network topology is quite common in networks of three nodes, appearing in around 18% of all network topologies included in this study (Figure 3E). To validate the importance of this structure, we computed the number of networks with an incoherent bivalent motif classified as hormetic by our analysis. Results (Figure 3F) show a 5X increase in the probability to exhibit hormesis, when comparing this subset (58% of the topologies with incoherent bivalent motif can show hormesis) with the set containing all topologies (11%, Figure 2A).

Next, we proceeded to analyze how extra links in the networks are capable of enhancing the rate of hormesis in a topology. To do this, we compare the percentage of biphasic dose-response curves for a given topology as we include more links (Figure 3G). Taking as a starting point the parental network of Group 3, we see how the new links around the incoherent bivalent motif have a mild positive effect on the rate of hormesis, increasing the number of parameter sets that show hormesis from 5% to 6%.

In conclusion, our analysis shows that hormesis requires a biphasic network motif centered on the *Target*. The two paths that act on the *Output* are required to be incoherent (when row one in the adjacency matrix is positive, row one is net negative (+,-), and vice versa). The hormetic topologies can be clustered in four families, corresponding to the four possibilities for this core motif. The probability of a biphasic dose-response curve in any of these four families can be increased by extra links to the network.

### The backward interaction needs to work in the saturated regime

Next, we focus on the dynamics of the interactions to search for relevant combinations of parameter values linked to biphasic dose-response curves. To do that, we combine in a violin plot (Figure 4A) all values of all kinetic and Michaelis-Menten constants that produced a biphasic drug response. Our analysis shows that most parameters produce hormesis in their entire range tested, with one exception, the Michaelis-Menten constant of the interaction from *Target* to *Input* K_21_. Apart from a small number of outliers, its value is significantly lower than the other Michaelis-Menten constants. From the point of view of the Michaelis-Menten interaction, this suggests that the Target is catalyzing the activation or inhibition of the *Input* node working in the saturated regime. In other words, the speed of this reaction is not dependent on the amount of substrate.

**Figure 4:**
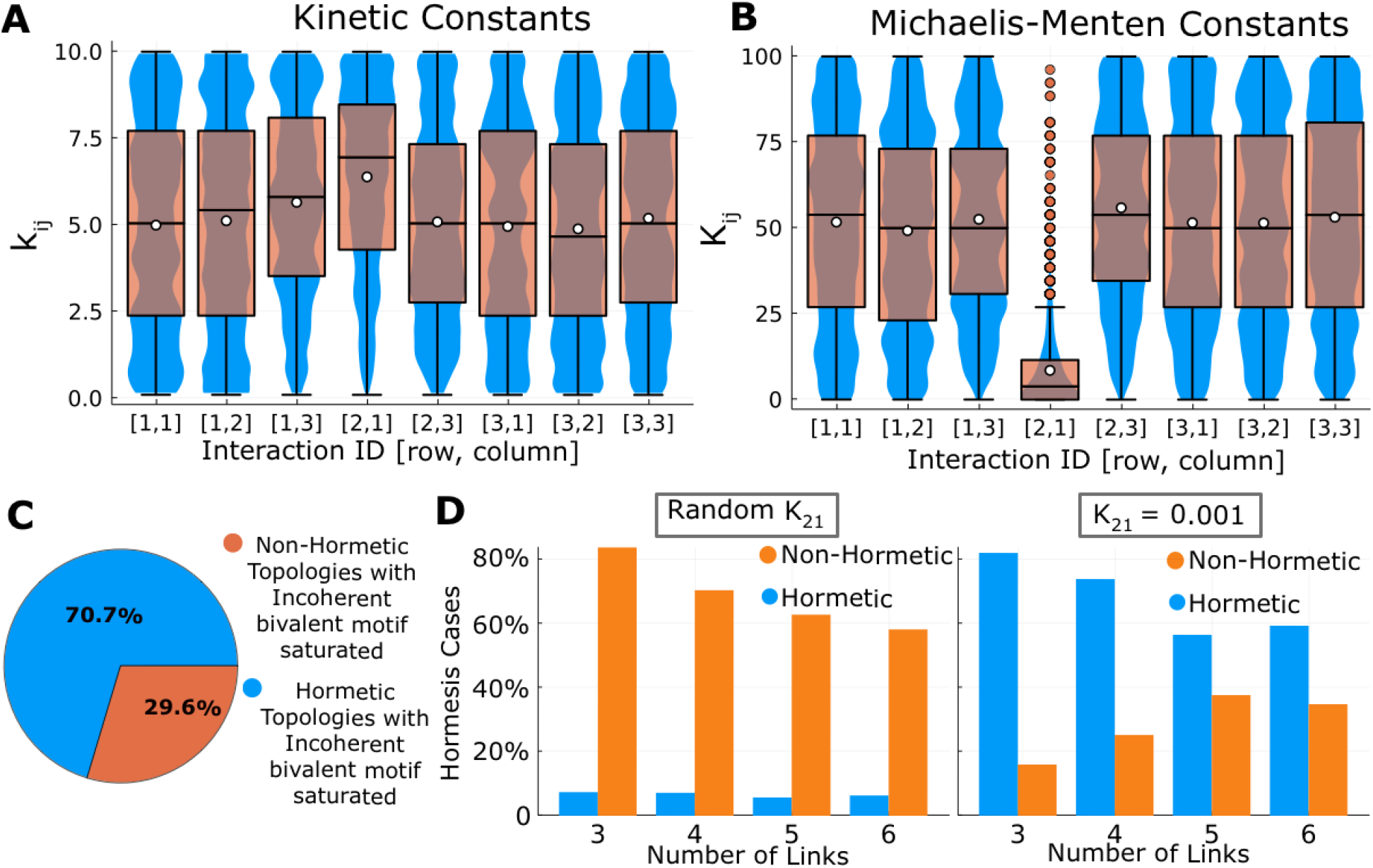
Hormesis requires the backwards reaction to work in the saturated enzymatic regime. (A-B) A violin diagram of all values of the kinetic constant and Michaelis-Menten constants that show hormesis. The blue region shows the distribution of values. The white dot is the median value. The orange dots are outliers (i.e., values outside the 1.5 × Inter Quartile Range, marked by the errorbars). (C) Distribution of hormetic and non-hormetic topologies that contain one of the Incoherent Bivalent motifs after fixing the value of K_21_ = 0.01 (to be compared with Figure 3F). (D) Comparison of the percentage of hormetic dose-response curves in the subset of topologies with Incoherent Bivalent Motifs when using a fixed K_21_ = 0.001 (green bars) versus randomly selected values (blue bars).

To study the impact of this low value of K_21_, we performed the analysis in all topologies that contain each of the possible four bivalent motifs (18% of the total topologies), but now fixing the value of K_21_ = 0.001. In these conditions, the percentage of topologies capable of producing hormesis increased from 58% (Figure 3F) to more than 70% (Figure 4C). Despite this mild increase, when focusing on the probability of hormesis for each topology, the true importance of K_21_ is shown (Figure 4D): from around 6% of parameter sets that produce hormesis (when K_21_ is random), the rate now increases to 80% when K_21_ is fixed at 0.001.

In conclusion, our analysis illustrates that the nature of the interaction between *Target* and *Input* has a clear effect on the probability of observing hormesis, with a 10x increase when the speed of this reaction becomes insensitive to the amount of *Input*. In other words, the combination of an incoherent bivalent loop centered in the inhibited node and the upstream reaction working in the saturated regime strongly increases the probability of producing hormetic dose-response curves.

### Rapamycin is hormetic due to a biphasic motif

As explained in the introduction, one of the most well-characterized and mediatic small molecule inhibitors is rapamycin, due to its potential in extending the life span of organisms. Many independent studies show a clear inverted U-Shape dose-response curve after long-term rapamycin inhibition (more than 24 hours of treatment), in terms of cell viability, mitochondrial density [23], and expression of Ki-67 (a marker for cell proliferation). The maximum efficiency takes place typically around 1 nM [42], with a clear reduction in these markers if rapamycin is applied at higher concentrations (up to 50 nM). Typically, studies attribute this reduction to side effects or cytotoxicity [43]. On the other hand, many studies often use rapamycin at 100 nM [44] and even 200 nM in tissue culture with no observable or measurable cytotoxic effects [45–50].

As an alternative hypothesis, we propose to investigate whether the interaction network in the mTOR-Pi3K pathway could be the cause of the hormesis for rapamycin. To test this hypothesis, we will follow a similar approach of computational high-throughput. Now, instead of the coarse-grained depiction of a pathway, we will use a full model that contains all the known interactions in the mTOR-Pi3K signaling [24], designed as a chain of kinase-phosphatase opposing pairs. The network of interactions is depicted in Figure 5A and explained in detail in the Methods section. In this scheme, the ligand upstream of the signaling cascade is insulin and insulin growth factors, and the downstream readout is identified as mTORC2, since it drives cell cycle progression and survival via Akt, also regulating mitochondrial physiology, metabolism, and integrity [51]. Moreover, mTORC2 activity has been shown to increase with age [52], and its elimination has been shown to alleviate aging progression [53].

**Figure 5:**
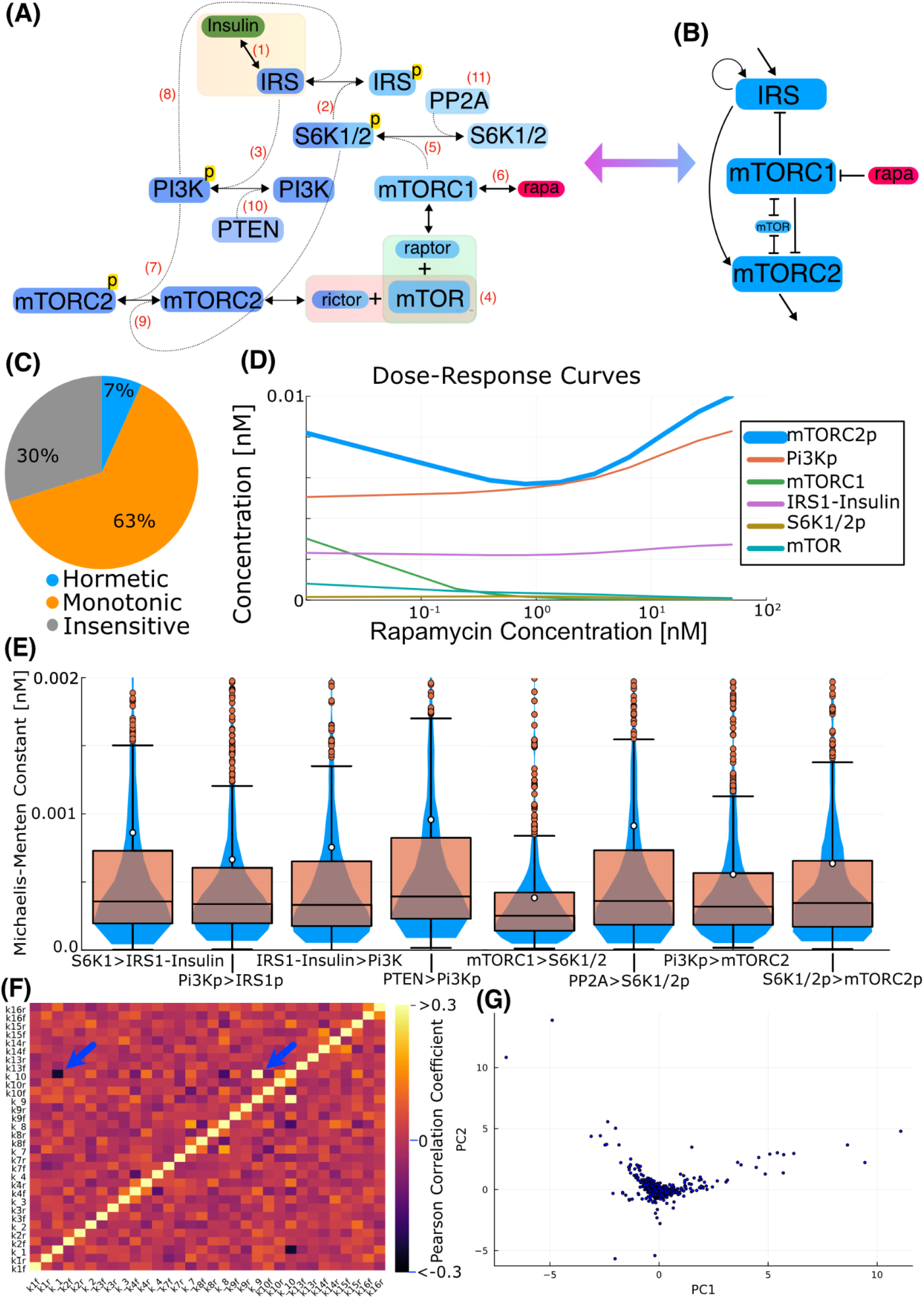
Rapamycin is Hormetic due to an incoherent bivalent motif centered in its target mTORC1. (A) Scheme of the interactions of the mTOR pathway included in the model. (B) simplification of the interactions in the mTOR pathway architecture as a 3-node network scheme to visualize the dual role of mTORC1. (C) Distribution of cases by shape of the Dose-response curve in the mTOR simulations. (D) Typical Dose-Response curve of active mTORC2 and other key molecules in the pathway. (E) Computation of the Michaelis-Menten constants in all catalyzed interactions involved in the signaling cascade. The blue region shows the distribution of values. The white dot is the median value. The orange dots are outliers (i.e., values outside the 1.5 × Inter Quartile Range, marked by the errorbars). (F) Heatmap plot of the Pearson Correlation coefficient between all parameters of the model. (G) Principal component analysis applied to the parameter sets that give Hormetic dose-response curves in the 16 core topologies. Higher coefficients in PC1 correspond to K_2_ and K_4_ (phosphorylation and dephosphorylation of Pi3K, respectively).

Following a similar strategy as in the previous sections (high-throughput numerical screening with parameter values selected from randomly distributed values), the mass action model composed of 25 coupled ODEs (see Methods) is solved for each parameter using different doses of rapamycin. The steady state value of the output (mTORC2 in this case) is used to generate dose-response curves that are then filtered using the same algorithm to account for bimodal shapes. Figure 5C resumes the results of this screening, showing that 70% of parameter sets in this topology show some sensitivity to rapamycin, with 10% of those dose-responses classified as biphasic. Taking into account that the phase space of the mTOR model is much larger than the coarse-grained model in terms of dimensions (24 parameters versus 16 for the 3-node model), this large probability of hormesis suggests that this topology is highly prone to exhibiting dose-response curves that are not the typical S-shape monotonic. Figure 5D illustrates a typical dose-response curve generated by the simulations, with a clear decrease in mTORC2 activity at intermediate levels (blue line), followed by a reduced sensitivity at higher doses. In addition, the model correctly reproduces the activation of Pi3K (orange line) after rapamycin treatment [54–55] and a reduction in the levels of free mTOR (cyan line) as the concentration of the inhibitor increases [56–57]. The model also reproduces the increase in activity of mTORC2 after short-term exposure to rapamycin [58]. On the other hand, long-term rapamycin exposure (around 24 hours) traps mTOR in mTORC1 complexes, and this obstructs the assembly of mTORC2 [59]. Therefore, rapamycin acts as an indirect activator of mTORC2 (via Pi3K) and an indirect inhibitor of mTORC2, resulting in a network structure that resembles the incoherent bivalent loop found to be at the core of biphasic dose-responses [60] (illustrated in Figure 5B).

To search for regions in the phase space where hormesis is more probable, we generate violin diagrams with all parameter values that produce biphasic dose-response curves. Results show that hormesis can occur in all parameter space for all parameters tested, but when we compute the Michaelis-Menten constants as K_i_ = (k_ir_+k__i_)/k_if_ for all interactions (Figure 5E), we observe that the constant corresponding to the activation of S6K1/2 by mTORC1 is significantly smaller in average, with a narrower distribution (blue regions), and lower medium (white dot). This correlates with our previous results that lower values of the Michaelis-Menten constant in the backwards interaction from the target (mTORC1 in this case) towards the Input (IRS1, controlled by S6K1/2 in this case) increase the rate of hormesis. Estimation of the Pearson correlation coefficient between all parameters of the model for the sets that produce hormesis is plotted as a heatmap in Figure 5F. This data shows a clear positive correlation between k__9_ and k__10_ (activation and deactivation of mTORC2). This is consistent with a required balance between opposite reactions acting in key nodes of the pathway. On the other hand, the analysis shows a significant negative correlation between k__1_ and k__10_ (inhibition of both IRS1-Insulin complex and mTORC2p by the same enzyme S6K1p). This correlation is interesting because both interactions negatively regulate mTORC2 at different levels of the cascade, so they should be negatively regulated to avoid excessive inhibition of mTORC2.

Next, to estimate the parameters that induce a higher effect in the hormesis, we reduced the dimensionality of the model using Principal Component Analysis (PCA), using all sets of Michaelis-Menten Constants that resulted in hormesis. The plot in Figure 5G shows all data represented in the obtained PC1 and PC2 axes. Interestingly, the most important parameters in PC1 are K_2_ = 0.66 and K_4_ = 0.7, which correspond to phosphorylation and dephosphorylation of Pi3K. This suggests that the levels of Pi3K (the kinase of our output mTORC2) are relevant in the mechanisms that induce the hormesis.

In conclusion, our analysis proposes an alternative explanation of the hormesis exhibited by rapamycin centered on the specific architecture of the network of interactions of the mTOR-Pi3K signaling cascade. This is potentially due to the presence of a topology that resembles an incoherent bivalent network motif centered in mTORC1 (illustrated in Figure 5B), which can induce a biphasic dose-response with high probability. The backwards reaction of this motif has a low Michaelis-Menten constant, consistent with our previous findings.

## Discussion

Biological systems process information through complex networks of biomolecular interactions that define how cells respond to stimuli and perturbations such as drug inhibition. Dissecting and simplifying these complicated networks in smaller, manageable key motifs and modules is a successful strategy to understand key properties of this signal processing [27, 61–64].

Previous theoretical approaches to studying hormesis focused on extracting curve parameters and on curve-fitting algorithms applied to these non-linear dose-responses [65–71]. More mechanistic approaches have also been developed to explain the hormesis in opiates based on the autocatalytic properties of opiate receptors [72] or the tradeoff of resources applied to chlorine as an oxidative stressor [73]. Here, we use an approach centered on a high-throughput computational screening that allows us to test a very large number of topologies and parameters [28, 74]. To our knowledge, our study constitutes the first attempt to search for general mechanisms that may induce complex non-monotonic responses.

Focusing on the limitations of our study, our coarse-grained perspective [75–77] of representing large pathways as simple networks of three nodes condenses molecular interactions upstream of a *Target* node as a single *Input* node, and molecular interactions downstream of our *Target* as a single *Output* node. This strategy implies that interactions upstream or downstream of the *Target* are linear or close to linear, and that nonlinear interactions are restricted as positive or negative feedback loops on *Input* and *Output* links, or between the three nodes (as Michaelis-Menten type). This simplification is required to obtain a manageable and simple-to-understand set of topologies that can be solved using a high-throughput type of approach. On the other hand, the purpose of the high-throughput approach is not to reproduce the behavior of a specific system, but to find general patterns and rules that can result in complex dose-response curves. In this direction, the fact that our approach allowed us to explain the tendency towards hormesis of a much more complex and realistic system (such as rapamicin) serves as a favorable argument about the usefulness of these types of coarse-grain approaches.

Another main limitation in our study is that it relies on a Michaelis-Menten simplification for the interactions between the nodes (implicitly assuming that enzyme-substrate complex dissociation is basically instantaneous compared to other processes, which may not apply to all steps of a signaling pathway. Alternatively, the model of *mTOR-Pi3K* has been written using a full mass action approach. Interestingly, the conclusions of the simplified 3-node Michaelis-Menten model translate quite well to the more complex full-mass-action approach. Based on this, we argue that the search for general rules using this type of simple conceptual models can be used to understand more complex and realistic systems.

In these scenarios of more complex models, still, some simplifications are required to make modeling approaches useful and not unnecessarily overly complicated. Our rapamycin model includes its binding to mTORC1, and a dynamic equilibrium between free mTOR and mTORC1 [12]. A more detailed model should include the direct intracellular receptor of rapamycin, FKBP12, which interacts with the FRB domain of mTOR. The RICTOR subunit of the mTORC2 inhibits its interaction with the FKBP12-rapamycin, making it effectively insensitive to rapamycin. Alternatively, our model includes the dynamic disassembly and turnover of the complex, mTORC2, and rapamycin sequestering in mTORC1 complexes, the mTOR required for mTORC2.

One of the main findings of our study is the role of the Michaelis-Menten constant of the backward activating or inhibiting interaction from the *Target* to the *Input*. This saturation acts as an effective non-linear threshold, i.e., the direct pathway to the Output is affected in a relatively linear manner by the inhibitor, but the indirect pathway is buffered (slight reductions in Target activity do not affect the Input node’s activity, and only take place once the inhibitor crosses a critical threshold. The fact that only the saturation of the backward link arises as relevant, suggests that the delayed response of the indirect path also plays an important role and contributes to the nonlinear nature of this link.

We believe that the main impact of our approach is that it elucidates how a network can shape and modify the response to a drug treatment by inducing complexity in the response. The two main ingredients unveiled by our study (an incoherent bivalent network motif centered on the target of inhibition, and the upstream interaction working in the saturated regime) seem to represent a general rule for a system to exhibit hormesis. A network structure similar to this motif is at the core of the *mTOR-Pi3K* pathway, and we propose that the hormesis of rapamycin may arise from this topology.

## Conclusions

Signaling pathways are highly nonlinear systems by design, and therefore, inhibitors of proteins inside these complex networks may exhibit a relation between dosage and response that is nonlinear. Therefore, as an alternative approach to discarding drugs from the drug development pipeline due to an atypical dose-response, we argue that a better strategy is to understand how this complex behavior arises and define an optimal dose or optimal strategy to overcome or even benefit from this nonlinear behavior.

Here, we performed an *in silico* screening of a coarse-grained model of a signaling pathway to study the minimal ingredients that will induce hormesis. Our analysis shows that complex dose-responses with biphasic shape arise due to a bivalent interaction centered at the target of inhibition, where the two interactions have an opposite effect (direct or indirect) over the output downstream (incoherent). In addition, if the upstream interaction is taking place in the saturated regime (in terms of a typical Michaelis-Menten approach), the resulting dose-response curves will be hormetic with much higher probability than just monotonic. This insight from abstract network topologies allowed us to understand that rapamycin may exhibit a biphasic dose-response due to a similar incoherent network motif around its target mTOR.

Overall, we argue that this type of simplified computational study can be very helpful to understand how complex systems such as signaling pathways integrate, interpret, and process signals, as well as to understand or even predict the effects and dynamics of drugs that target elements inside a highly nonlinear network architecture.

## Notes

https://github.com/GuillermoCerrillo/TFM_Hormesis

### Competing Interest Statement

The authors have declared no competing interest.

### Funding

- Ministerio de Ciencia e Innovación, Spain: RTI2018-096953-B-I00, PID2022-140421NB-I00, PDC2022-133147-I00, CEX2023-001316-M

